# A novel method to enhance the sensitivity of marker detection using a refined hierarchical prior of tissue similarities

**DOI:** 10.1101/020685

**Authors:** Shahin Mohammadi, Ananth Grama

**Affiliations:** Department of Computer Sciences, Purdue University, West Lafayette, IN, USA

## Abstract

**Motivation:** Identification of biochemical processes that drive the transformation of a totipotent cell into various cell types is essential to our understanding of living systems. This complex machinery determines how tissues differ in terms of their anatomy, physiology, morphology, and, more importantly, how various cellular control mechanisms contribute to the observed similarities/ differences. Tissue-selective genes orchestrate various aspects of cellular machinery in different tissues, and are known to be implicated in a number of tissue-specific pathologies.

**Results:** We propose a novel statistical approach that identifies and removes the effect of universally expressed genes in groups of tissues. This allows us to better characterize tissue similarities, as well as to identify tissue-selective genes. We use our method to construct a reliable hierarchy of tissue similarities. The groupings of tissues in this hierarchy are used to specify successively refined priors for identifying tissue-selective functions and their corresponding genes in the reduced subspace. We show that our refinement process enhances the signal-to-noise ratio in the identification of markers. Using case studies of immune cells and brain tissues, we show that our approach significantly outperforms the state-of-the-art methods, both in terms of coverage and reliability of the predicted tissue-selective genes.

**Conclusions:** Our statistical approach provides a general framework for enhancing the sensitivity of marker detection methods, which can be used in conjunction with other techniques. Even in cases where the number of available expression datasets is limited, we show that our marker detection method outperforms existing techniques. We present detailed validation on immune cells and brain tissues in this paper. Our approach can be applied to construct similar datasets of other human tissues as well, for identifying tissue-specific genes. We demonstrate how these tissue-selective genes enhance our understanding of differentiating biochemical features of brain tissues, shed light on how tissue-selective pathologies progress, and help us identify specific biomarkers and targets for future interventions.

## Background

An embryonic stem cell contains all genetic information needed to develop a new individual; it differentiates into various cell types, which group together to shape tissues, combine to constitute organs, and assemble into organ systems. Uncovering these sophisticated processes is integral to our understanding of tissue ontogenesis. Various differentiated tissues, while inheriting a similar genetic code, exhibit unique anatomical and physiological properties. These spatio-temporally differentiated characteristics are achieved through systematic control of cellular machinery at different levels, including transcriptional, translational, and post-translational regulations, in order to orchestrate tissue-specific functions and dynamic responses to environmental stimuli.

Transcriptional regulation is a well-studied aspect of this control. It is manifested in the observed differences in expression levels of genes across tissues. *Housekeeping genes* are universally expressed in human tissues and undertake core cellular functions[1, 2, 3]. In contrast, certain genes are specifically or preferentially expressed in one, or a set of biologically relevant tissue types[4, 5, 2, 6]. *Tissue-selective genes* are highly under/over expressed in a selected set of tissues, as compared to other tissues. *Tissue-specific genes* constitute a special case of tissue-selective genes, in which a target gene is uniquely expressed in a single tissue type, but not in other tissues. Interestingly, a number of known disease genes are tissue-selective and are under/over expressed in the specific tissue(s) where the gene defect causes pathology[7, 8]. To this end, tissue-selective genes play a crucial role in the physiology and the pathophysiology of human tissues. They are not only critical factors in elucidating the molecular mechanisms of normal tissues, but are also promising drug targets and candidate biomarkers.

Various methods have been proposed in literature to identify and characterize genes that are preferentially expressed in different tissues. These methods can be broadly classified into two groups: i) methods that use a single expression profile of each tissue/ cell-type; and ii) meta-analysis approaches that collect and use multiple biological instances of each tissue/ cell-type into a comprehensive profile. Schug *et al.*[32] proposed an information theoretic approach that is an instance of the former class. They used Shannon’s entropy to measure uniformity of expression across tissues and rank genes based on their overall tissue-specificity. Kadota *et al.*[9] proposed a variation of the entropy-based method, called ROKU, which pre-processes expression profiles using Tukey biweights to allow identification of both up/ down regulated genes. More recently, Cavalli *et al.*[5] proposed a new method, called SpeCond, which models the background expression of each gene using a Gaussian mixture model. It then assigns a *p*-value to the unusually high/low expression values in specific tissues, relative to this null distribution.

Meta-analysis approaches, on the other hand, use a comprehensive expression profile aggregated over different studies. These methods use multiple biological replica of each tissue/ cell-type to distinguish between biological and technical variation of genes. Wang *et al.*[4] present among the first methods in this class. They manually annotated different studies available in the NCBI Gene Expression Omnibus (GEO) database and compiled a comprehensive dataset of tissue-specific expression profiles. For each group of tissues, they used dChip to normalize the raw expression profiles and correct for technical variation using a model-based approach. Using this aggregated profile, they employed an equal weighting strategy among expression profiles to identify tissue-selective genes. Teng *et al.*[6] used the same input dataset, but applied a supervised learning scheme, based on support vector machines (SVM) and random forests (RF), to automatically assign a reliability weight to each expression profile with respect to a manually curated set of true positives collected from Uniprot database.

These methods have contributed significantly to the understanding of tissue-specific processes. In this paper, we aim to further build upon these efforts with the goal of enhancing their performance. Our work is premised on the following observations: First, there is a significantly-expressed set of common housekeeping functions across human tissues, which potentially masks less-expressed tissue-specific pathways, making it harder to identify preferential markers. We hypothesize that by deflating known prior signal contributed by housekeeping genes, one can enhance the signal-to-noise (SNR) ratio of the tissue-specific expression profiles. This improves the sensitivity of prediction methods for identifying cell-type markers. Second, the rationale behind the concept of tissue restriction is that genes that are expressed in more tissues are related to generic cellular functions, whereas genes that are expressed in only a few selected tissued are involved in specific functions and are more informative regarding the biological processes that uniquely differentiate tissues from each other. While this concept is intuitive, the use of frequency/ tissue-counts is not an appropriate measure of specificity. The underlying similarity of tissues in which the gene is active also plays a key role in defining its specificity. For example, a gene that is expressed in a large number of lymphocytes supports specificity much more so than a gene that is expressed in the same number of biologically unrelated tissues. Finally, in cases where the number of available replicas are limited, biological variation of expression is hard to estimate, which can affect the reproducibility of the results.

Motivated by these considerations, we propose a novel marker detection method, that significantly enhances the sensitivity of previous methods in identifying tissue-selective/ specific expression, as well as in approximating the biological variation of expression, given a limited set of expression profiles. To this end, we first organize a given set of tissues and cell-types into a hierarchy such that tissues at the same level exhibit similar levels of functional similarity. We use singular value decomposition (SVD) to estimate the common signature of genes, and couple it with a statistical approach to adjust the *raw transcriptional signatures* for the effect of housekeeping genes. We then identify groups of similar tissues using these adjusted signatures, and contract these similar groups to construct a reduced space, which enhances the inter group separability. Finally, we present a novel method that utilizes this refined hierarchical prior, constructed as the reduced subspace over groups of similar tissues, and identifies genes that are preferentially expressed within each group at each level of the hierarchy.

Using known immune-cell markers, we show that our *adjusted transcriptional signatures* enhance the mean expression level of the marker genes in their host cell-type, defined as the *signal*, while both decreasing their expression level in unrelated cell-types, and silencing the expression of housekeeping genes, defined as *noise* in this context. This supports our hypothesis that eliminating the common signature of housekeeping genes enhances the signal-to-noise (SNR) ratio for identifying marker genes. Moreover, we show that adjusted transcriptional similarities, computed as the the normalized dot-product of the adjusted transcriptional similarities, can distinguish both similar tissues and dissimilar pairs of tissues.

We apply our method to identify novel brain-selective genes, and cross-validate our results with a recently published brain-specific proteome[11]. Compared to sets of related genes from prior studies, our method has higher sensitivity in extracting brain-enriched genes, especially in cases where only a limited number of expression profiles per tissue/ cell-type are available. Most human pathologies are driven by a combined perturbation of universal, housekeeping genes coupled with tissue-selective elements that make the target tissue uniquely susceptible to the disease. These tissue-selective genes provide ideal candidates as disease biomarkers, as well as, drug targets, since they have the potential to directly affect a selected set of tissues, without disrupting the normal functions of other tissue types.

## Results and discussion

### Adjusting raw transcriptional signatures enhances signal-to-noise (SNR) ratio

Housekeeping (HK) genes are ubiquitously expressed across all human tissues/ cell-types, and are responsible for maintaining core cellular functions, including but not limited to translation, RNA processing, intracellular transport, and energy metabolism[1, 2]. While computational methods for identifying housekeeping genes differ in their underlying assumptions[12], they share the same goal – identifying molecular targets, such as genes and their corresponding proteins, which are needed by all cell types in the body. These universal genes define a shared functional space across human cells, which is also known to be highly conserved across species. However, while fundamental for our understanding of cell biology, these genes do not provide insights into the unique characteristics of different tissues. When comparing different tissues to identify their similarities and differences, these highly expressed genes may dominate differentiating signal [13].

To address this problem, we propose a method based on a singular value decomposition (SVD) to estimate the expression profiles of housekeeping genes. Using this housekeeping signature, we adjust raw transcriptional signatures of human tissues/ cell-types by applying our proposed Peeling Common Signature (PCS) method to silence the shared housekeeping prior (please refer to Materials and Methods section for more details). The adjustment procedure is shown in Figure 1, and both the raw and adjusted transcriptional signatures are available for download as Additional file 1 and Additional file 2, respectively. To assess the quality of adjustment procedure, we perform two independent experiments using housekeeping genes and known marker genes.

**Figure 1:**
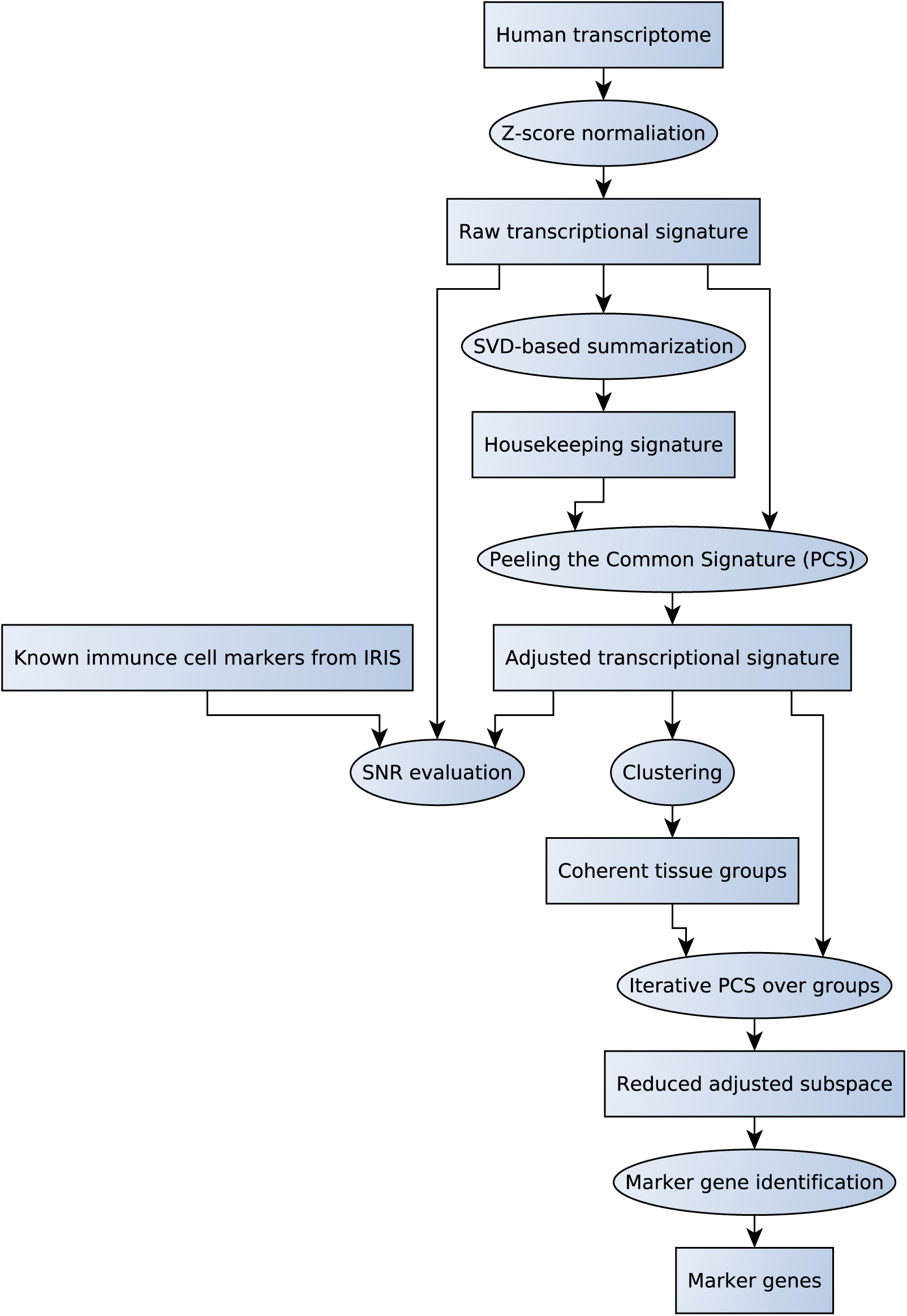
Overall workflow of our method. Transcriptional signatures are normalized, before/after deflating housekeeping signatures, and used for marker detection

First, we evaluate whether our adjustment method indeed deflates the effect of housekeeping genes. To this end, we investigate two well-studied housekeeping genes – *Glyceraldehyde 3-phosphate dehydrogenase (GAPDH)* and *β-actin* for further analysis. These genes are routinely used as control probes, including in the Affymetrix microarray platforms. For each of these genes, we average all the probesets that are mapped to them, and plot the mean value across all tissues in this study. Figure 2 (a) and Figure 2 (b) show the effect of this adjustment on the two genes. For *β-actin*, we observe a uniformly high expression across all tissues/ cell-types before the adjustment (median before adjustment is 4.83). After adjustment, we observe a significant deflation in the plot (median after adjustment is -0.05). On the other hand, *GAPDH* seems to have additional tissue-specific roles in muscle and brain tissues. Its gene product encodes one of the 10 key enzymes in the glycolytic pathway, and is previously known to be expressed in higher-levels in energy-demanding tissues[14]. Nevertheless, this gene is expressed at high levels in all human tissues (median before adjustment is 4.08), which is reduced after adjustment (median after adjustment is -0.26). Additionally, to assess if a similar pattern emerges if we use multiple housekeeping genes, we downloaded the selected set of 11 high quality housekeeping genes, other than GAPDH or *β*-actin, from a recent study[12]. Figure 3 illustrates the average of these control housekeeping genes across different tissues, before and after adjustment. Unlike GAPDH, these probes seem to be uniformly high in most tissues, though lower than GAPDH and *β* actin on average, but are under-expressed in a small number of tissues. Regardless, we observe a similar deflation pattern from a median of 1.82 to 0.05 after adjustment. This supports the hypothesis that our adjustment process correctly deflates the expression level of housekeeping genes in the adjusted signatures.

**Figure 2:**
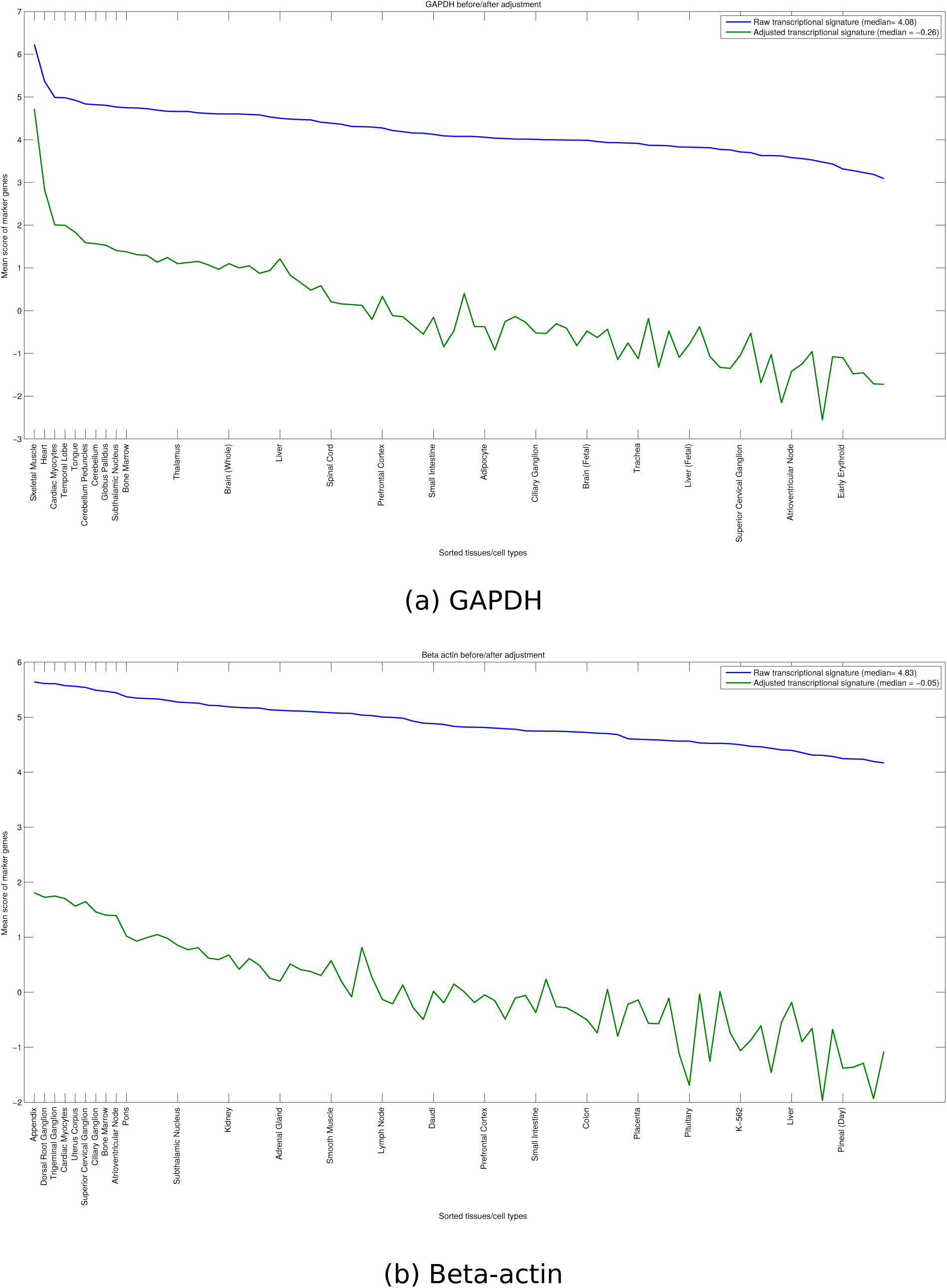
Effect of adjustment on the expression level of housekeeping genes. Affymetrix probes sponding to GAPDH and *β*-actin are averaged independently for raw and adjusted transcriptional signatures, and tissues are sorted based on their mean raw values.

**Figure 3:**
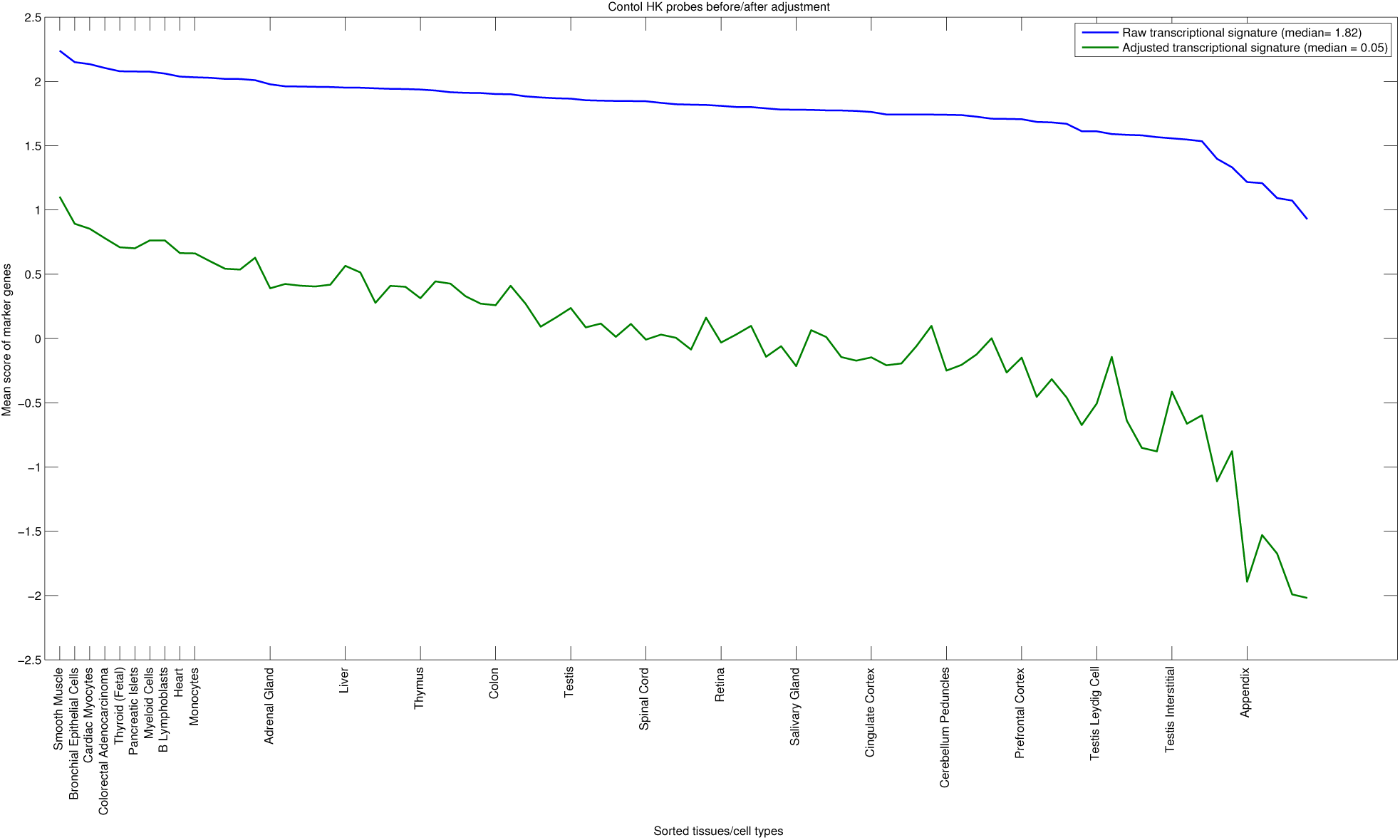
Effect of adjustment on the expression level of control housekeeping genes. High quality control probes from Eisenberg and Levanon[12] were used to assess the performance of adjustment method on reducing the noise contributed by housekeeping genes

Next, we investigated the effect of transcriptional adjustment on marker genes. Tissue-selective genes, while highly informative, are expressed at lower levels compared to housekeeping genes. By suppressing the “noise” level contributed by housekeeping genes, transcriptional adjustment boosts the SNR and sensitivity, which in turn allows us to identify more reliable cell markers.

Immune cells are among the most studied cell-types in human body, and a number of high-quality markers are known for major immune cell classes. To assess our hypothesis, we adopted the gold-standard of immune-cell specific markers from the Immune Response In Silico (IRIS) dataset[15], which is based on a meta-analysis of multiple microarray expression profiles for six key immune cell types. We mapped probesets in this dataset to the probesets in the GNF Gene atlas, and manually matched the studied cell-types in IRIS to the immune cell-types in our dataset. Among these cell-types, we removed ones that are not matched. Furthermore, we observed that the dendritic cell (DC) markers in this dataset are myeloid-derived DC cells, whereas the DC cell profiled in GNF gene atlas is plasmacytoid DC cell (from lymphoid lineage). These two type of cells are distinct from each other, and none of the markers were applicable here, so we also filtered DC from further study.

Table 1 summarizes the statistics of all selected cell-types and their corresponding markers. We study each of these cell-types individually to better understand the effect of adjustment on increasing the *signal* level. It is notable here that the average value of markers identifies both the signal, in the related tissues, and the noise, in unrelated tissues. Figure 4 illustrates the effect of adjustment on three major lymphoid-derived immune cells, namely B-cell, T-cell, and NK-cell. B-cell and T-cell markers behave similarly, that is, the mean value of marker signal is increased in the relevant cell-types, and reduced in the rest of tissues/ cell-types. For the adjusted profile of B-cell, the B-cell itself has the highest expression level for these markers, followed by DC cell, tonsil, and Daudi cell-line. Plasmacytoid DC cells express CD123, CD303, and CD304 surface markers, but lack CD11c (mDC marker) and CD14 (monocyte marker). Unlike conventional DC cells, which are monocyte-like, pDC cells are known to exhibit a high similarity to plasma B-cells [16]. Tonsil is also a rich source of B-cells exhibiting a variety of phenotypes and activation states, and have been used routinely to isolate highly purified tonsil B lymphocytes[17]. The Daudi cell-line is a well characterized B-lymphoblast cell-line, isolated from peripheral blood, which has been used extensively to study Burkitt’s lymphoma. These tissues/ cell-types serve as the “host” for B-cell markers, and the average expression value of these markers in the host cells signifies the signal level. As clear from Figure 4 (a), the average value of markers has been increased in the adjusted signatures compared to the raw signatures. Moreover, the expression of B-cell markers in unrelated tissues on right-side of the plot is considered noise, and reduced by our method. Similarly, the most significant hosts of T-cell are thymus (in which T-cells mature), MOLT-4 (which is a T lymphoblast cell-line), and T-cells (both CD8 (killer) and CD4 (helper) cells). Average expression levels of these T-cell markers have also increased in the relevant hosts, and decreased otherwise. Finally, we notice that for NK cell markers, there is a significant drop after the first cell-type (NK cell). This signifies the specificity of NK cell markers, or it can be an artifact of the lower number of available markers to assess for NK cell.

**Table 1:** Sumary of the immune cell markers from the IRIS dataset

| Cell-type | # IRIS probes | # matched probes | # unique genes |
| --- | --- | --- | --- |
| B Cell | 121 | 66 | 54 |
| Dendritic Cell | 86 | 48 | 41 |
| Monocyte | 103 | 67 | 53 |
| Myeloid | 449 | 246 | 201 |
| NK Cell | 24 | 18 | 15 |
| T Cell | 94 | 55 | 46 |

**Figure 4:**
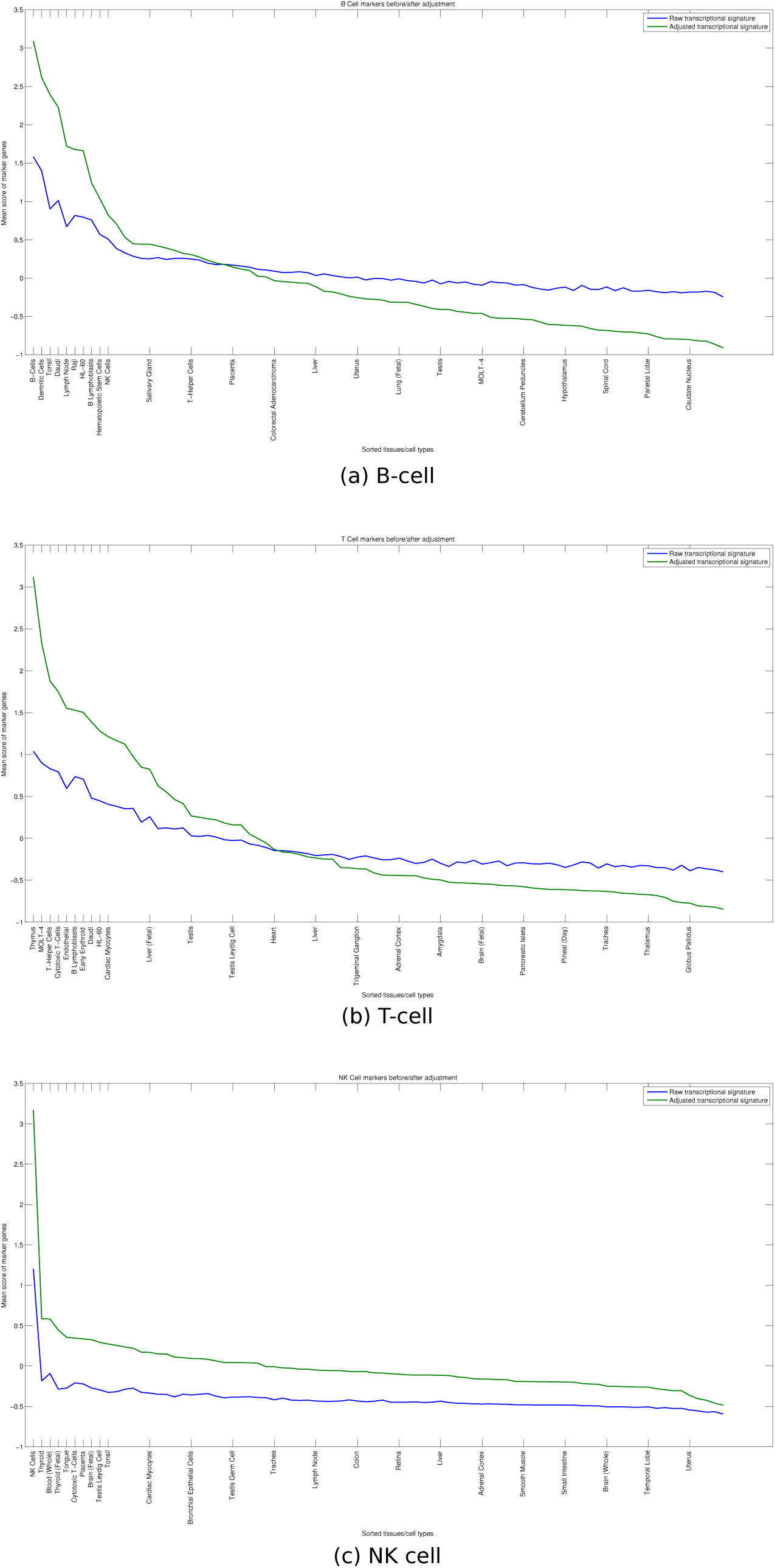
Effect of adjustment on the expression level of marker genes. Each sub-plot corresponds to one of the major sub-types of lymphoid-derived immune cells. Tissues and cell-types are sorted based on the average value of markers in the adjusted signature profile.

Figure 5 illustrates the effect of adjustment on markers of the fully differentiated monocytes, as well as the myeloid-progenitor cells isolated using CD33 marker. Both of these plots exhibit similar characteristics to the B-cell and T-cell plots– in the sense that the signal level has been elavated in the relavant host cells and decreased in unrelated tisssues/ cell-types. Monocytes are derived from the myeloid-lineage, and as such, share a number of markers with their progenitor cell. This can be verified by the existence of both monocyte and myeloid cells in the host cells of both plots. However, myeloid cells also differentiate into a variety of other cell-types, including platelets, granulocytes, and erythrocytes (red blood cells), all of which are circulating in the blood stream. This has been illustrated by having whole blood as the top host for myeloid cells, but having lower-rank for monocytes.

**Figure 5:**
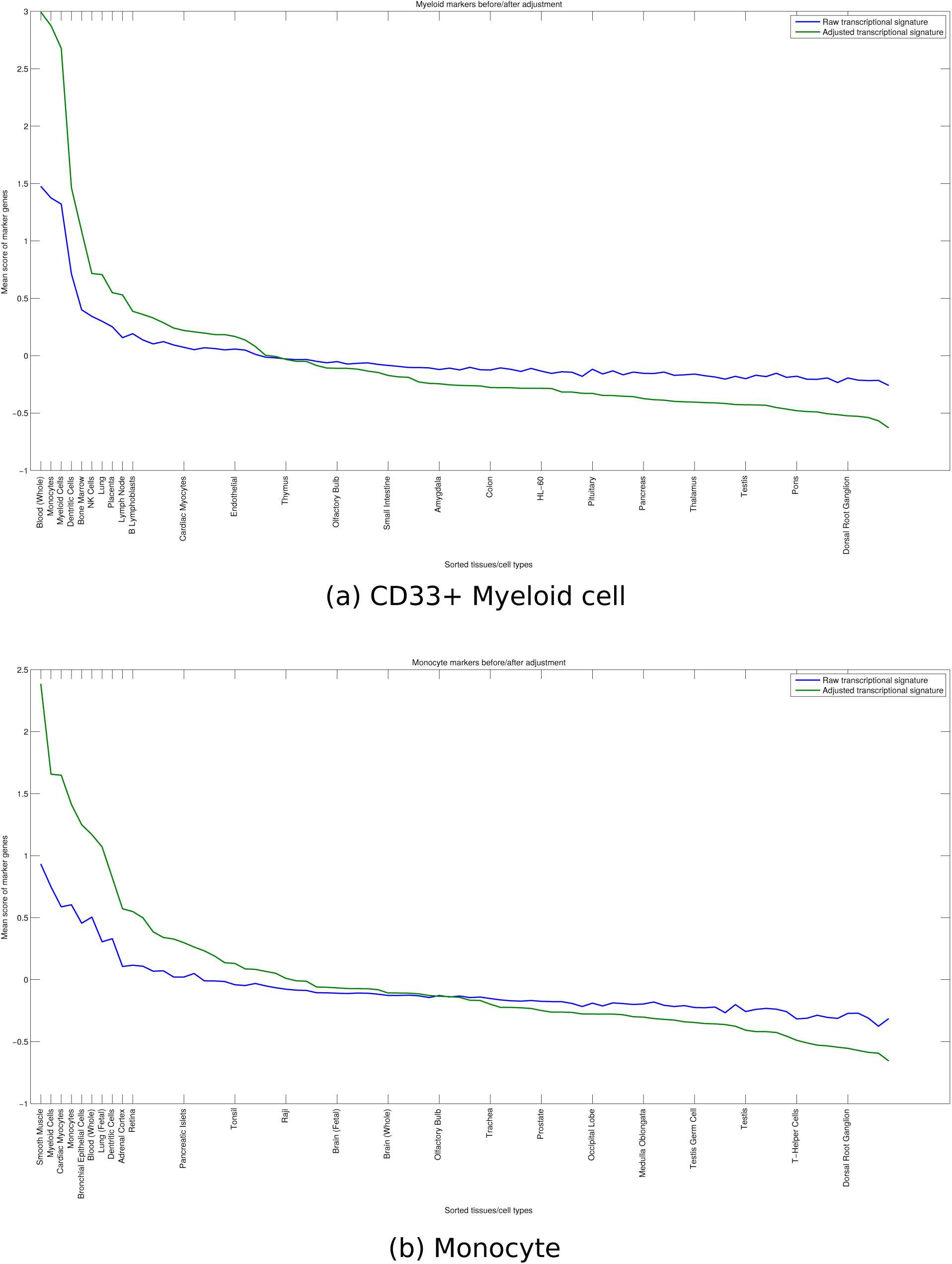
Effect of adjustment on the expression level of marker genes. Each sub-plot corresponds to one of the myeloid immune cells. Tissues and cell-types are sorted based on the average value of markers in the adjusted signature profile.

Another interesting observation is that pDC cells, while derived from lymphoid lineage, share markers with the myeloid cells. More interestingly, two involuntary muscles, namely the smooth muscle and cardiac myocyte (heart muscle), are among the highest-ranked hosts for monocyte markers. In order to verify the monocyte markers that have higher signal in these muscle tissues than in monocyte itself, we extracted shared markers that show higher expression level in both of these tissues. This results in 35 probesets corresponding to 30 unique genes, the list of which is available for download as Additional file 3. We used g:Profiler[18] to assess the functional enrichment of this ordered list of genes. We focused on GO biological processes, as well as KEGG and Reactome pathways, and used a *p*-value cutoff of 1*e* – 3 after correcting for multiple hypothesis testing using Benjamini-Hochberg method. We used moderate hierarchical filtering to select best GO terms per parent group. The sorted list of enriched functional terms is available for download as Additional file 4.

Among these terms, there are two major groups: (i) genes that are involved in *inflammatory response*, *cytokine-cytokine receptor interaction*, and *positive regulation of cell migration*, and (ii) genes involved in extracellular matrix (ECM) remodeling, or more specifically *Collagen degradation*. The former set is enriched in cytokines including *CXCL1/3/5*, *IL1A/B*, and *IL6*, whereas the later group is enriched in the matrix metalloproteases (MMPs), such as *MMP1/9/14/19*. These genes are zinc-dependent endopeptidases that are the major proteases involved in regulating matrix remodeling component of the MMP cluster. To further investigate if similar pattern emerges in meta-analysis of enriched genes using a compendium of cell-type specific microarray datasets, we also constructed the EnrichmentMap of monocyte markers using the Gene Enrichment Profiler[19]. Among these genes, we manually highlighted shared markers that show high enrichment in smooth muscle and cardiac myocyte to investigate shared markers.

The final highlighted enrichment map is available for download as Additional file 5. Among shared genes in this map, *PTX3, TFPI2, MMP1,* and *MMP14* are highly enriched in both smooth muscle and cardiac myocyte, whereas *PLAUR, IL6, IL1A, IL1B, CXCL1, CXCL3, ATP13A3, CXCL5, PID1, IER3,* and *VCAN* are enriched in smooth muscle. This suggests that the ECM remodeling pathways are utilized by both involuntary muscles, while inflammatory response pathway is mostly enriched in smooth muscle. There is support in literature for the role of inflammation and cell migration in smooth muscle cells [20, 21, 22, 23, 24, 25], as well as extracellular matrix remodeling [26, 27]. Our results suggest a greater role for shared inter-cellular signaling pathways, which merits further investigation.

### Pairwise transcriptional similarities after adjustment differentiate similar versus dissimilar tissues

Ubiquitous expression of housekeeping genes in different tissues, in turn, contributes to a significantly high transcriptional correlation between all pairs of human tissues. This baseline similarity makes it hard to distinguish between similar, unrelated, or dissimilar tissues. To remedy this problem, we propose an alternate statistical framework based on partial Pearson’s correlation, or equivalently the normalized dot-product of adjusted transcriptional signatures, to estimate the overall similarity of tissues/ cell types while controlling for the effect of housekeeping genes. A similar approach has been previously applied to the problem of gene regulatory network (GRN) inference to distinguish direct versus indirect interactions between pairs of related genes [28].

Figure 6 shows the distribution of pairwise transcriptional similarities between every pair of tissues/ cell-types, before and after adjusting for the effect of housekeeping genes. Raw transcriptional similarities are inflated by the presence of housekeeping genes, and have a tight peak around its mean of 0.83, which makes discriminating similar/ dissimilar tissues hard. On the other hand, the mean of adjusted similarity scores is close to zero (*-* 0.01), which suggests that majority of tissues/ cell-types are unrelated after removing the known prior contributed by housekeeping genes. The distribution of adjusted similarity scores, however, is not symmetric around the mean and has longer tail for the positive similarities, which indicates that the maximum similarity among tissues is much higher than the maximum dissimilarity. This dissimilarity can be interpreted as an orthogonality in the functional space of the corresponding tissues/ cell-types, that is, they use complementary sets of pathways. Adjusted similarity scores below the 10^*th*^ percentile and above the 90^*th*^ percentile are color-coded by red and green, which mark the space of dissimilar and similar tissues/ cell-types, respectively. The complete table of adjusted scores is available for download as Additional file 6.

**Figure 6:**
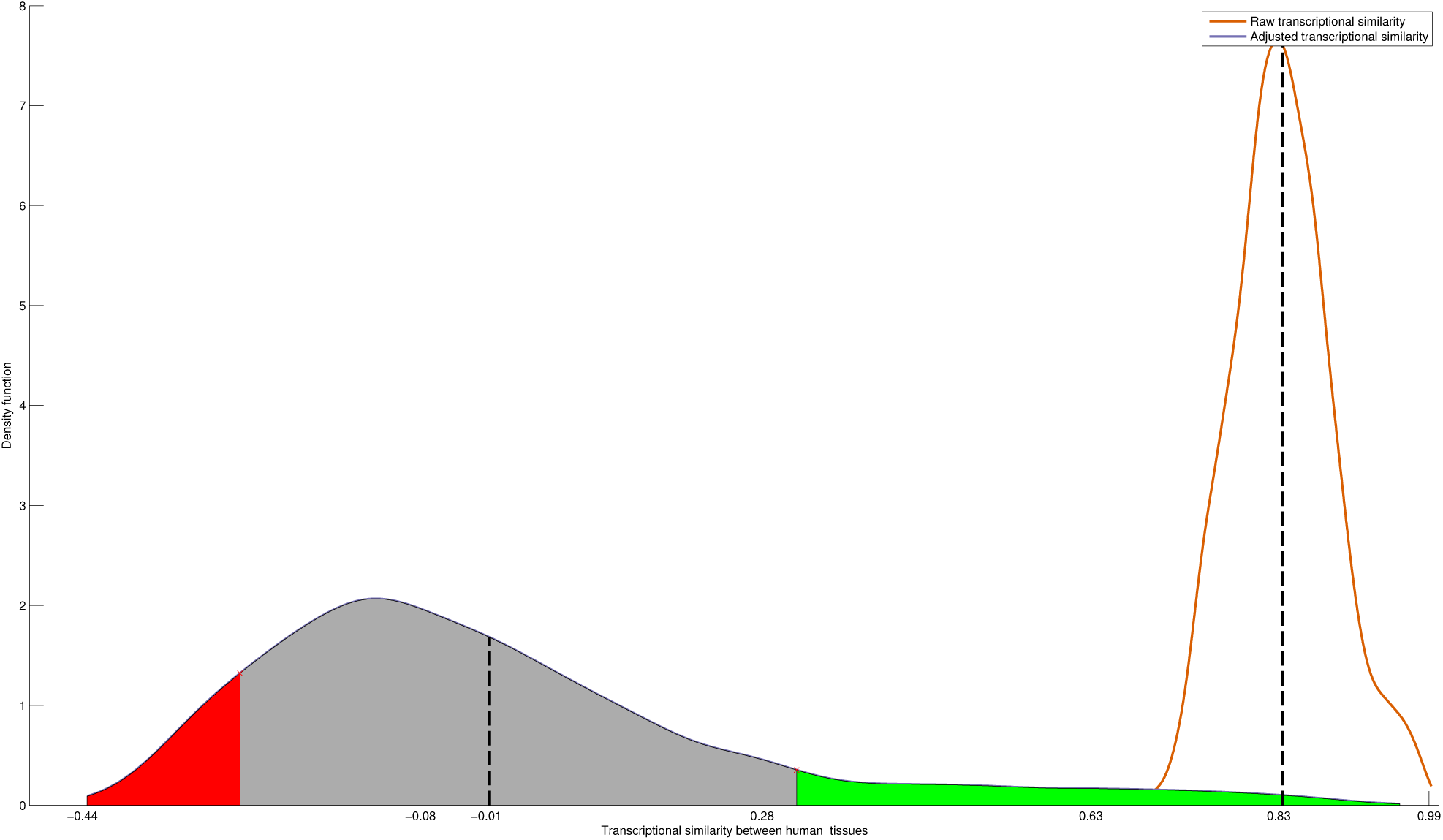
Distribution of tissue-tissue similarities before/after adjustment. Similarity scores are computed using Pearson’s correlation and partial Pearson’s correlation, or equivalently, the normalized dot product of raw versus adjusted transcriptional signatures. Raw transcriptional similarities are artificially inflated due to the present of housekeeping genes, which makes discriminating similar/dissimilar tissues hard. On the other hand, adjusted similarities are centered around zero, which suggests that majority of tissues/ cell-types are unrelated after removing the known prior contributed by housekeeping genes.

To better understand how different tissues/ cell-types interact with respect to their similarity/ dissimilarity scores, we constructed a pruned Tissue-Tissue Similarity Network (TTSN) by setting a stringent threshold and extracting all pair-wise interactions that are lower than the 1^*st*^ percentile or greater than the 99^*th*^ percentile. We used Cytoscape[29] to import and visualize this network, which is shown in Figure 7. A visual inspection of the map reveals multiple groupings of similar tissues, including core brain tissues and testis tissues, as well as similar pairs including cerebellum tissues, pineal gland, and immune cells including myeloid-derived cells and T-cells.

**Figure 7:**
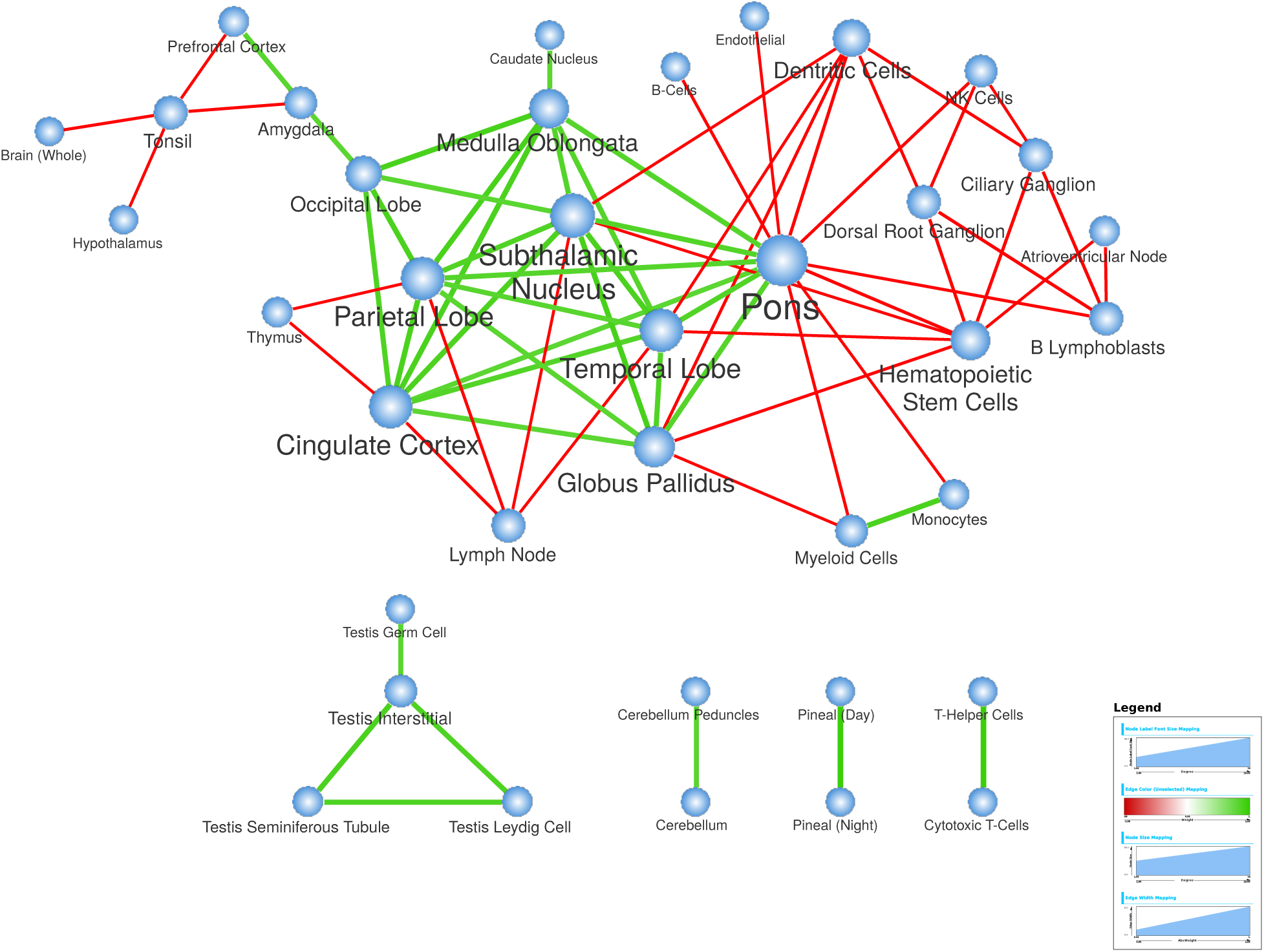
Network of most significant pairwise relationships among human tissues/ cell-type. Each node in this network represents a human tissue or cell-type, and edges represent the most significant pairwise relationships among them. Red edges correspond to dissimilar pairs, while green edges correspond to similar pairs. The color gradient of edges is used to represent the strength of the relationship, whereas node sizes correspond to the degree of each node.

Additionally, there are reoccurring patterns of dissimilarity between immune and neural cells. One such example is reflected in the enriched negative edges between tissues involved in the maturation and storage of lymphocytes, namely tonsil, thymus, and lymph node, and tissue samples from different regions of the brain. Another example is the pons tissue, which acts as a negative hub, showing dissimilarity to a wide range of immune cells, Finally, we observe that ganglion-related tissues, while acting as part of the peripheral nervous system, do not show significant similarity with the brain tissue, but in the negative space of dissimilarities, they exhibit similar negative pattern with immune cells. This suggests that the negative space of dissimilarities contains information, much like the positive space of similarities regarding the utilization of functional pathways between pairs of human tissues/ cell-types. This is masked if the effect of housekeeping genes is not suitably deflated.

### Tissue-tissue similarities cluster into coherent groups of similar tissues

As observed in the investigation of pair-wise tissue similarities in the previous section, human tissues fall within dense groups that exhibit high degree of connectivity within the cluster, while having clear separation from the rest of tissues. To identify such major groups, we cluster adjusted transcriptional signatures using agglomerative hierarchical clustering. We use 1-adjusted transcriptional similarities (partial Pearson’s correlation between pairs of tissues after correcting for the common HK signature) as the distance measure between tissues, and the unweighted average distance (UPGMA) method for computing the distance between clusters. To cut the clustering tree into distinct clusters, we set a threshold of 0.5 on the average distance between clusters. This results in 39 clusters, corresponding to 16 non-trivial clusters, which we manually label using the dominant pattern of member tissues in each cluster. We observe a similar pattern, which is tight grouping among brain tissues and immune cells, respectively, and significant dissimilarity between these clusters. To further analyze this pattern, we use the identified core brain cluster and the immune cell cluster as centers and sort other clusters based their relative distance to these two poles. Figure 8 shows the computed heatmap over non-trivial clusters, after sorting and annotating each cluster accordingly. Interestingly, three major groupings appear in among clusters: (i) neural cells involved in both central and peripheral nervous system; (ii) glandular tissues including both tissues residing in the brain as well as other anatomical locations, and (iii) immune cells and cell-lines. Among glandular tissues, pineal gland shows the highest similarity with the brain cluster, while adrenal gland does not. Interestingly, involuntary muscles, including smooth muscle and cardiac myocyte, group closely with immune cells, whereas skeletal muscle shows higher similarity with the nerve cells in PNS. This is also consistent with the observed overlap among the monocyte markers and involuntary muscles, which we noted in previous sections. Finally, we note that there is a significant dissimilarity between immune cell clusters and both CNS an PNS, which is also consistent with our previous observation over the tissue-tissue similarity network. The complete list of non-trivial clusters is available for download as Additional file 7. We use these groupings as a refined prior to contract the space of adjusted transcriptional signatures before identifying tissue-selective genes in order to enhance our ability to assess context-restriction of markers.

**Figure 8:**
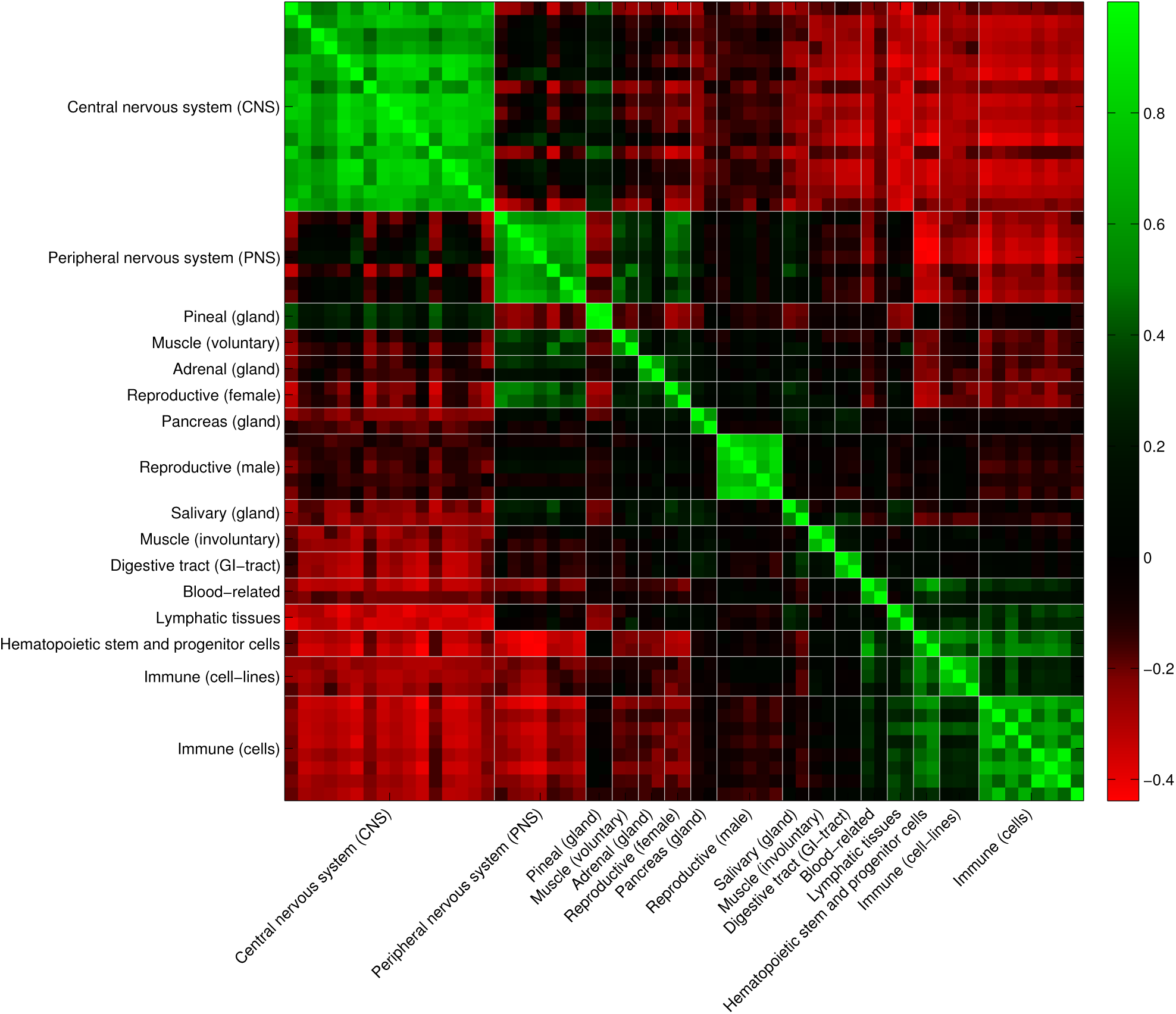
Heatmap of cluster tissues according to adjusted transcriptional similarities. Each cluster is annotated using the majority of its member tissues and visually separated from the rest of clusters using vertical/horizontal separators. From top to bottom (or left to right) there are three major groups of tissues: (i) brain tissues, (ii) glandular tissues, and (iii) immune tissues/ cell-types

### Reducing the space of adjusted tissue signatures provides a refined prior for identifying marker genes

Tissue-specific genes, which are expressed in only one specific tissue/ cell-type, represent the most conservative subset of marker genes. Depending on the structure of input tissues/ cell-types, it can be difficult to find such genes if there is a high-level of dependence/ correlation among different columns of the signature matrix. Moreover, many tissue-selective genes have preferential expression pattern in multiple tissues to perform either similar or completely different roles. GNF gene atlas contains tissues, cell-types, and transformed cell-lines that have a considerable overlap with each other. Previous studies, including the one conducted by Cavalli *et al.*[5], manually select tissues to avoid this problem. However, this has two main drawbacks: (i) the process needs manual intervention and understanding of the underlying similarity of given tissues/ cell-types, which can be biased, based on the functional aspect under study; and (ii) there is a significant loss of information regarding genes that are consistently used in similar tissues. Specially in cases in which there is a lack of biological replicas to estimate biological variation of genes, such as the case with GNF gene atlas, this will prevent us from having an estimate of the expression variance. To remedy these problems, we propose to use the same SVD-based method, described earlier for identifying HK signature, and apply it independently to each identified cluster and represent each cluster using its unique signature. Using this approach, we reduced the whole matrix of adjusted transcriptional signatures among 84 tissues/ celltypes, into a projected space of 39 signatures (16 approximated signatures for each non-trivial cluster, plus the adjusted transcriptional signature of singleton clusters). The reduced matrix of adjusted signatures is available for download as Additional file 8. We will use this matrix for all our further analysis to find marker genes.

### Using the reduced subspace of adjusted signatures enhances the sensitivity of marker detection methods– case study of brain tissue markers

#### Identifying brain markers and brain-specific functional pathways

To demonstrate the effectiveness of the reduced subspace of adjusted signatures in identifying makers, we present a case study of applying our new method to identify novel brain-selective markers. We chose brain tissue for further investigation because it has the highest number of curated profiles in the meta-analysis methods, covering more than 20% of all aggregated microarrays. The rest of tissues had significantly lower coverage.

For setting the parameters in our method, we assume equal variance in the active and null populations, and set a *p*-value threshold of 1*e* – 5 to declare that genes exhibit selective expression domain. In terms of tissue-restriction, we limit markers to the ones that are expressed in at most half of the tissues under study. Finally, we set a *separability gap* of 0.25 to ensure that activated population has a uniformly high expression value. The complete list of all identified markers for different tissue groups, including markers in central nervous system (CNS), is available for download as Additional file 9.

To identify functional pathways corresponding to the identified brain markers, we use g:Profiler[18] to find enriched terms in Gene Ontology (GO) biological processes, KEGG and Reactome pathways, and the Human Phenotype Ontology (HPO). We set a *p*-value cutoff threshold of 1*e* – 3, after correcting for multiple hypothesis testing using Benjamini-Hochberg method, to declare a term as significant. In order to simplify the list of enriched term using the underlying DAG structure of GO and HPO, we use a moderate hierarchical filtering to select the best terms per parent group. The complete list of enriched functional terms for brain marker genes is available for download as Additional file 10. The top 20 enriched terms are presented in Table 2. These terms cover major brain functions, from low-level cellular functions, such as synaptic transmission, glutamate secretion, and axon guidance, to high-level behavioral terms, such as learning, memory, and morphine addiction. Interestingly, these enriched terms also include brain-specific disorders, such as epileptic encephalopathy. This is consistent with our prior understanding– that is, while most human pathologies are driven by a combined disregulation of universal, housekeeping functions coupled with tissue-selective elements, the tissue-selective component plays a major role in unique susceptibility of host tissue to the tissue-specific disorders[7, 8]. For some brain-specific disorders, such as neurodegenerative disorders ranging from Parkinson’s, Alzheimer’s, and Huntington’s disease, the housekeeping element, such as the role of oxidative phosphorylation, is better understood. However, our results suggest that for many other brain-related disorders, such as epilepsy and schizophrenia, brain-specific markers play a crucial role. We expand on this claim in the following subsections.

**Table 2:**
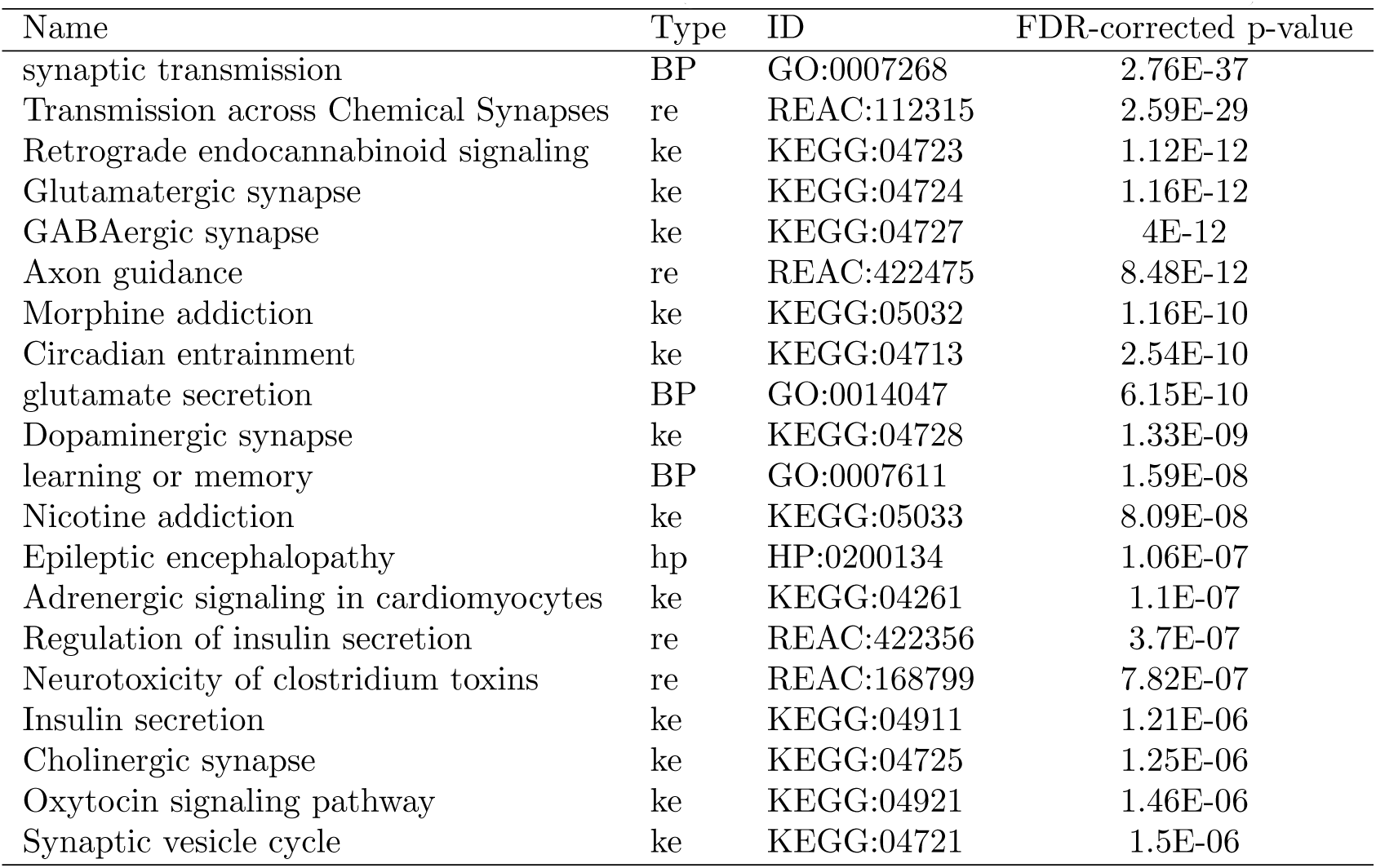
Top 20 enriched terms among identified brain markers in this study

#### Measuring the extent of overlap in brain-selective genesets from different studies

To evaluate the extent of overlap among the previously reported brain-specific genes, we extracted and consolidated four different gene-sets from prior studies. Cavalli *et al.*[5] *fits normal mixture model with three components to the expression profile of each gene in a selected subset of 32/ 79 tissues from GNF Gene Atlas. Then, they identify the null distribution* among these fits and use it to identify outlier conditions/ tissues for each gene. We collected 511 up-regulated genes in the *whole brain* tissue as candidate brain-specific genes in this study. Then, we mapped the Ensembl gene identifiers in this dataset to their corresponding Hugo gene symbols using Ensembl BioMart[31], resulting in 497 unique genes. Next, we extracted marker genes from the study conducted by Schug *et al.*[32], *which uses an entropy-based method and is the ancestor of ROKU method[9]. This method, unlike ROKU, focuses on the up-regulated genes in different tissues. We follow the same protocol proposed by the authors and use a cutoff threshold of less than or equal to 7 on the Q* value (categorical selectivity) of RMA normalized probesets in the ”cortex,” which results in 708 brain-selective probesets corresponding to 628 genes. Finally, we compared our method to two other studies that are based on meta-analysis of the microarray profiles. These methods use a compendium of tissue/ cell-type specific microarray datasets, instead of a single study. Wang *et al.*[4] and Teng et al.[6] both use the same input dataset for their analysis, namely 616 brain-specific gene expression profiles based on the Affymetrix HG-U133 Plus 2.0, but follow a different strategy to assign weights to each chip and select tissue-selective genes. We gathered 222 and 1,408 brain-specific probesets from these studies and used Ensembl BioMart to convert these probeset ids to their corresponding HUGO gene symbol, resulting in 152 and 848 unique genes, respectively. The final set of brain-markers in these studies mapped to their gene symbol is available for download as Additional file 11.

To qualitatively assess the overlap among these sets, we construct the Venn diagram of intersections among these datasets, which is illustrated in Figure 9. A visual inspection of this diagram suggests that these methods cover vastly different sets of the brain-selective genes, with only 17 genes being in common among all methods. Next, to quantify pairwise overlaps among these datasets, we use the Jaccard index. Given a pair of marker sets, *A* and *B*, we calculate the Jaccard index is follows:

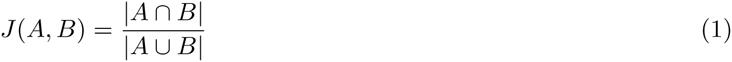

**Figure 9:**
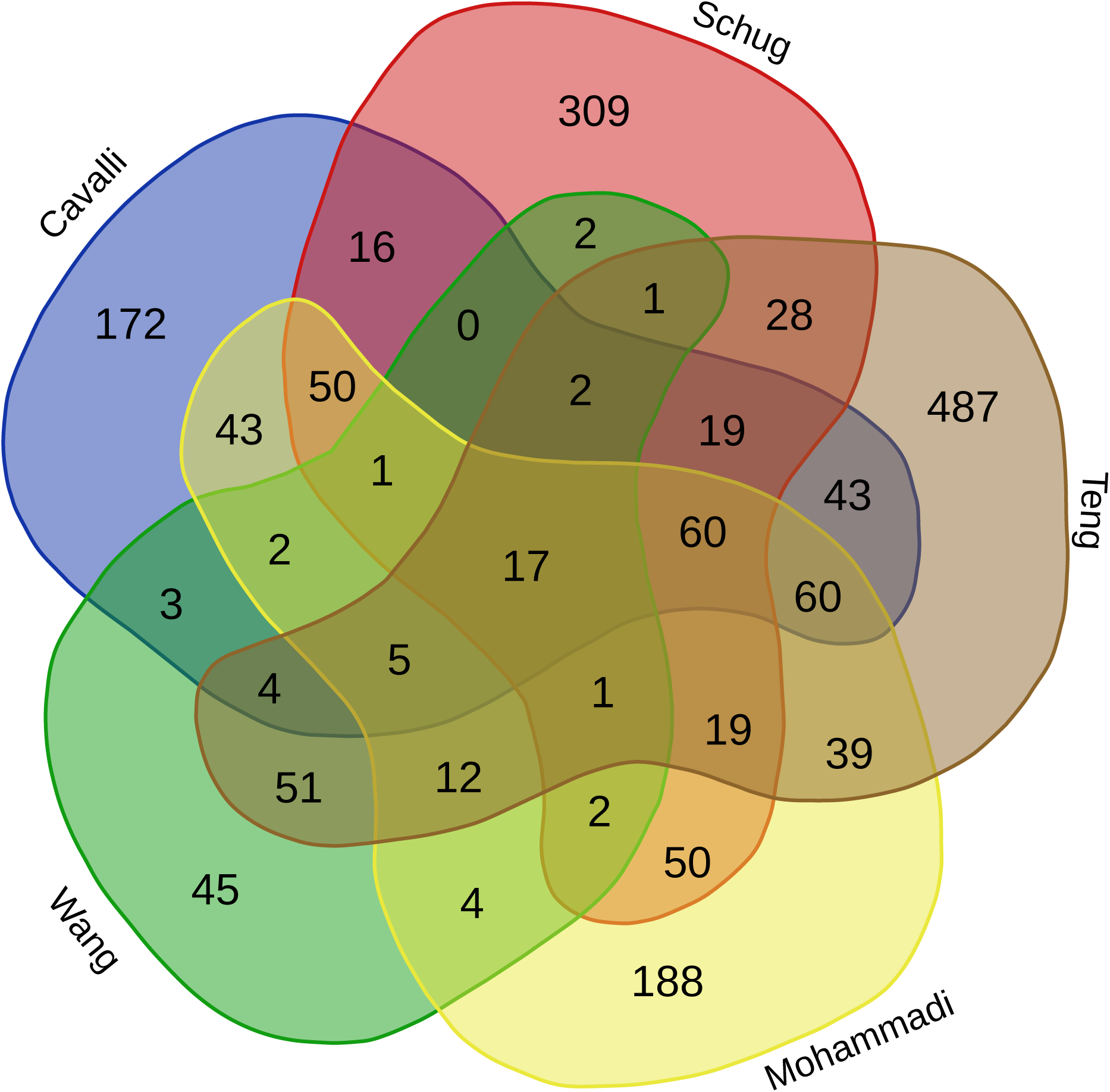
Venn diagram of the overlaps among brain-specific genesets. Each dataset is represented by a separate closed curve, named by its first, and the overlap set are color-coded independently and annotated by the number of genes that reside in each set

Table 3 summarizes the pairwise overlap among different datasets. Wang *et al.* dataset shows the least overlap with other datasets, with the greatest overlap being with the Teng *et al.* dataset. This is partially expected since both of these methods use the same aggregated input panel. Our dataset, on the other hand, shows the highest overall overlap with other methods, except with the Wang *et al.* dataset. Indeed, this can be observed by noting the fact that except for the last row (which corresponds to Wang *et al.* dataset), the maximum value of overlap is always on the second column (corresponding to our dataset). This suggests that our method combines the characteristics of both class of methods, which in turn allows us to achieve high sensitivity even when only a limited number of input datasets are available.

**Table 3:** Extent of overlap among brain marker genesets from different studies, measured using Jaccard index

|  | Cavalli | Mohammadi | Schug | Teng | Wang |
| --- | --- | --- | --- | --- | --- |
| Cavalli | 1 | 0.31234 | 0.22949 | 0.21388 | 0.06038 |
| Mohammadi | 0.31234 | 1 | 0.25316 | 0.22187 | 0.07051 |
| Schug | 0.22949 | 0.25316 | 1 | 0.16154 | 0.04943 |
| Teng | 0.21388 | 0.22187 | 0.16154 | 1 | 0.11027 |
| Wang | 0.06038 | 0.07051 | 0.04943 | 0.11027 | 1 |

#### Validating brain-specific genes in different studies using the gold standard from the human brain-specific proteome

In order to validate our predicted brain markers, and compare our method with previous studies, we use a recently published map of the human tissue proteome, which is based on combined transcriptomics profiling, using RNASeq, coupled with an antibody-based proteomics assay, using microarray-based immunohistochemistry[11]. The *brain-specific proteome* in this study contains 1134 genes whose expression is elevated in the brain, among which 318 are uniquely enriched in brain. The concept of brain-elevated versus enriched corresponds to our definition of tissue-selective and tissue-specific, respectively. We used the Human Protein Atlas[30] portal to download brain-elevated genes, identified using the RNASeq dataset, which are also supported by the immunohistochemistry experiments. This results in 273 total genes that are supported by both experiments, which map to 246 genes that are assayed in our study. We will use this set of genes as the gold standard to evaluate different brain marker genesets.

We measure various statistics for each dataset. First, we set the 17,220 genes assayed by GNF as the global sampling population, and map all marker sets to this *universal set*. Then, we computed the true-positives as the number of genes in each marker set that overlaps with the gold standard from the human brain-specific proteome. We approximated true negatives as the number of genes that are neither in the marker geneset nor the brain-specific proteome. Finally, we compute sensitivity, specificity, accuracy, and the hypergeometric *p*-value for each dataset. The final set of computed statistics is presented in Table 4. Please note that the computed specificity and accuracy are not exact, since the assumption that any gene not identified by the brain-specific proteome is a true negative, is not valid. This dataset is a high quality gold standard set, which only reports a gene as positive if there is a strong evidence for it. In other words, it has much higher chance of having false negatives than false positives. The hypergeometric *p*-value is the most statistically sound measure to compare these methods. As evident from the results, our method significantly outperforms prior methods that use a limited number of expression profiles, while only trailing the Teng *et al.* dataset, which uses a comprehensive aggregated dataset of brain-specific panel as its input. This suggests that by leveraging information across different tissues/ cell-types, our method can estimate the biological variation of selective genes, even in cases where multiple biological replicates are not available.

**Table 4:** Evaluation of brain marker genesets from different studies using the brain-specific proteome

|  | # markers | # mapped markers | True-positives | Sensitivity | Specificity | Accuracy | p-value |
| --- | --- | --- | --- | --- | --- | --- | --- |
| Cavalli | 497 | 447 | 88 | 0.358 | 0.979 | 0.970 | 3E-77 |
| Mohammadi | 553 | 553 | 105 | 0.427 | 0.974 | 0.966 | 6E-92 |
| Schug | 628 | 437 | 80 | 0.325 | 0.979 | 0.970 | 5E-67 |
| Teng | 848 | 620 | 120 | 0.488 | 0.971 | 0.964 | 5E-108 |
| Wang | 152 | 115 | 30 | 0.122 | 0.995 | 0.983 | 1E-29 |

#### Power of brain-selective genes in characterizing brain-specific disorders

To evaluate the power of marker genes as biomarkers for tissue-specific disorders in brain, we used the Disease Ontology Semantic and Enrichment analysis (DOSE)[33] R/ Bioconductor package to assess the enrichment of the Disease Ontology (DO)[34] terms among brain-specific genes. We limit the terms to the ones that have at least five member genes, and use a *q*-value cutoff of 0.05 to filter enriched terms after FDR-correction. The final table of the terms that are enriched by at least one of the methods is available for download as Additional file 12. This dataset includes 52 terms, 18 of which are identified by all methods. These shared, enriched terms are summarized in Table 5. As a general pattern, all genesets are enriched with genes involved in brain-specific disorders, ranging from cognitive and psychotic disorders to neurodegenerative diseases. Similar to our validation results using Human Protein Atlas (HPA), the dataset of Wang *et al.* performs poorly, whereas Teng *et al.* dataset outperforms other methods in a majority of cases. While makers from most methods are highly enriched in more generic terms, each method outperforms in specific subsets of disorders. For example, the dataset of Schug *et al.* is significantly enriched for a host of disorders involving cognitive and psychotic disorders. Surprisingly, for cognitive disorders, dementia, Alzheimer’s disease, and tauopathy, Schug *et al.* dataset of brain-specific genes outperforms all other datasets, including the HPA gold standard set.

**Table 5:** Disease enrichment of brain-specific markers

| Disorders | Cavalli:qval | HPA:qval | Mohammadi:qval | Schug:qval | Teng:qval | Wang:qval |
| --- | --- | --- | --- | --- | --- | --- |
| disease of mental health | 1.01E-23 | 1.59E-33 | 1.85E-24 | 1.90E-32 | 4.43E-34 | 2.01E-14 |
| schizophrenia | 9.33E-21 | 8.76E-32 | 1.34E-23 | 3.06E-30 | 3.20E-23 | 5.52E-15 |
| psychotic disorder | 9.33E-21 | 8.76E-32 | 1.34E-23 | 3.30E-30 | 3.20E-23 | 5.52E-15 |
| cognitive disorder | 1.06E-20 | 8.70E-28 | 1.31E-22 | 1.25E-31 | 6.36E-23 | 5.52E-15 |
| brain disease | 9.61E-09 | 1.30E-14 | 1.45E-14 | 1.76E-13 | 3.59E-19 | 3.22E-07 |
| epilepsy syndrome | 1.11E-04 | 6.65E-10 | 5.27E-10 | 4.48E-10 | 1.02E-18 | 5.93E-07 |
| pervasive developmental disorder | 4.42E-06 | 1.71E-07 | 9.06E-09 | 2.02E-03 | 6.09E-18 | 1.55E-04 |
| autism spectrum disorder | 1.54E-05 | 3.27E-07 | 2.98E-09 | 1.16E-03 | 2.42E-17 | 1.13E-04 |
| autistic disorder | 1.54E-05 | 3.27E-07 | 2.98E-09 | 1.16E-03 | 2.42E-17 | 1.13E-04 |
| central nervous system disease | 4.88E-13 | 1.05E-12 | 3.98E-17 | 4.15E-13 | 5.84E-17 | 3.28E-07 |
| developmental disorder of mental health | 5.31E-06 | 6.88E-12 | 1.02E-08 | 3.98E-07 | 9.20E-17 | 3.90E-05 |
| neurodegenerative disease | 1.49E-08 | 3.27E-07 | 2.24E-10 | 3.29E-06 | 2.82E-05 | 5.81E-07 |
| mood disorder | 1.14E-03 | 1.51E-06 | 1.75E-04 | 1.25E-05 | 3.13E-09 | 3.99E-07 |
| bipolar disorder | 1.22E-03 | 1.87E-05 | 4.15E-04 | 3.05E-06 | 3.13E-09 | 8.91E-08 |
| nervous system disease | 7.50E-06 | 1.58E-06 | 3.89E-09 | 1.07E-08 | 5.42E-09 | 2.49E-04 |
| dementia | 8.30E-07 | 7.51E-05 | 4.76E-07 | 1.70E-07 | 1.99E-03 | 1.07E-04 |
| Alzheimer's disease | 1.40E-05 | 2.14E-03 | 1.78E-04 | 3.45E-06 | 3.27E-03 | 4.09E-04 |
| tauopathy | 1.54E-05 | 2.29E-03 | 2.03E-04 | 4.02E-06 | 3.68E-03 | 4.17E-04 |

This suggests that different methods, while having a considerable overlap, identify unique subsets that play key roles in different pathways involved in brain-specific pathologies. This is consistent with our results in previous sections showing that these methods provide complementary sets of marker genes, which merits further investigation on developing integrative methods to combine these results.

## Conclusion

A key challenge in identifying tissue-selective genes is to identify the set of tissues in which they are manifested. This task is further complicated by the underlying similarity of tissues/ cell-types, which imposes a natural grouping among them. In this paper, we presented a novel statistical framework for constructing a refined hierarchical prior of tissue similarities to organize them into a well-separated, reduced subspace. This organization is constructed from the raw expression profiles by first deflating the shared functionality attributed to housekeeping genes. Then, we identified group of similar tissues in this adjusted space, and contracted each tissue group by iterative application of our SVD-based method to identify the common signature of each cluster. Finally, we proposed a new method to utilize this reduced sub-space as a refined prior to identify genes that are preferentially expressed at each level of the hierarchy.

Using a case study of immune-specific and brain-related genes, we have shown that the set of identified marker genes in this study significantly overlaps with the most recent gold standard markers. These findings greatly enhance our understanding of tissue-selective functions beyond the set of “first-order” related tissues in a way that has not been possible though individual investigation of human tissues.

## Materials and Methods

### Datasets

#### Tissue-specific mRNA expression profiles

We downloaded the pre-processed Genomics Institute of the Novartis Research Foundation (GNF) Gene Atlas[35] dataset from the BioGPS portal[36, 37]. Using a combination of the Affymetrix HG-U133A and a custom-designed (GNF1H) array, a total of 84 unique human tissues and cell-types from different donors have been profiled in this study, which together interrogate 44,775 human transcripts covering both known, as well as predicted and poorly characterized genes. The raw intensity values have been normalized using GCRMA and averaged across all replicas. We use the log2-transformed expression value as a proxy for the gene activity throughout our study.

#### Probeset-to-gene annotations

We downloaded the HG-U133A annotations in *\*.chip* format from the GSEA FTP website. This dataset consists of annotations for 22,283 probesets in the Human Genome U133A chip. We downloaded Human GNF1H chip annotation, update May 2007, from the BioGPS portal[36, 37]. This table includes a total of 22,492 unique probesets for GNF1H, as well as 66 duplicate targets from U133A for the normalization procedure. We extracted the HUGO gene symbols for each of the unique probesets in this table and created the corresponding *\*.chip* file for GNF1H. Finally, we merged the annotations for both arrays to create a unified annotation table for the Gene Atlas, which contains 44,775 unique probesets.

We further filtered this dataset to remove Affymetrix control probes, ambiguous probes (mapped to more than one gene), and unmapped probes (probes that do not match any gene). This results in a total of 28,649 probesets, which are mapped to 17,220 unique gene symbols.

### Identifying the shared subspace among a group of tissues

A given set of tissues/ cell-types typically share a common set of genes/ pathways, while specializing through preferential genes that control and regulate this core shared set. Let us represent the *raw transcriptional signature* of these tissues using a matrix *T*, in which rows correspond to genes and columns correspond to different tissues. We are interested in finding the subspace of common genes, and to use it to adjust these transcriptional signatures. When the given set includes all, or majority of, human cell-types, the shared signature represents the signature of housekeeping genes.

There are a number of methods for approximating this common signature in *T*, the simplest of which would be to find the mean of its columns. A more elaborate approach involves decomposing *T* into sum of rank-one matrices, using methods such as *singular value decomposition (SVD)* or *non-negative matrix under-approximation (NMU)*. The general goal of these methods is to represent *T* as a sum of outer products of vectors. More formally, we write *T* as follows:

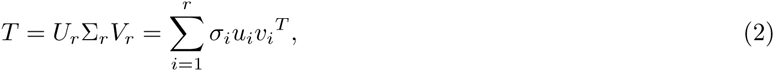

where *r ≤* min(*n*_*A*_, *n*_*B*_) is the rank of the approximation. In the SVD formulation, *u*_*i*_ and *v*_*i*_ vectors are called left and right singular vectors, respectively. These vectors constitute an orthonormal basis, that is both 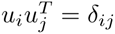 and 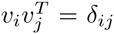 for all *i* and *j*. Additionally, for any *r*, an SVD is the optimal rank-*r* approximation of *T*. When all entries of *T* are positive, Perron-Frobenius theorem ensures that all entries of the both left and right singular vectors are positive. However, the first residual matrix, *R*_1_ = *M* – *σ*_1_*u*_1_*v*_1_^*T*^, can, and typically does, contain negative elements to ensure orthonormality. On the other hand, *NMU* formulation does not ensure orthonormality, but, rather enforces an additional constraint on the optimization problem, which is that *R*_*k*_ should consist of all positive elements. Unfortunately, while SVD has an optimal solution, the additional non-negativity constraint of NMU makes its computation non-convex, though heuristic exists to approximate the solution.

In this work, we use a rank-one approximation of matrix *T*, that is *r* = 1, to identify a unique signature that closely represents the common signature in *T*. We use the first singular vector of matrix *T*, after *z*-score normalization, as a proxy for the housekeeping signature throughout our study.

### Adjusting the raw transcriptional signatures to control for the known prior similarity

Let us denote the transcriptional profile of the *i*^*th*^ tissue by **T**_*i*_. In order to compute the *raw transcriptional similarity* between each given pair of tissues, we compute the Pearson’s correlation as follows:

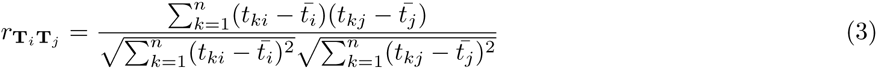

in which *n* represents the total number of genes, *t*_*ki*_ and *t*_*kj*_ are the expression levels of the *k*^*th*^ genes in the *i*^*th*^ and *j*^*th*^ tissues, respectively. Similarly, 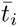 and 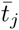 represent the average expression level of genes in the corresponding tissues. Let **Z**_*i*_ denote the *Z*-score normalized version of **T**_*i*_, defined as 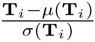. We will refer to **Z**_*i*_ as the *raw transcriptional signature* of tissue *i*. Using this formulation, we can simplify the raw transcriptional similarity as the normalized dot product of raw transcriptional signatures:

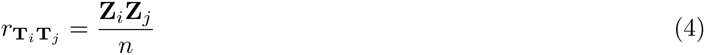

The raw transcriptional similarity of tissues is artificially inflated due to the ubiquitous expression of house-keeping genes across all tissues. To control for this effect, we first define the *housekeeping transcriptional signature*, denoted by vector **S**, as the left singular vector of matrix **Z**. Using this notation, we revise our similarity scores by computing the partial Pearson’s correlation between **T**_**i**_ and **T**_**i**_, after controlling for the effect of **S** as follows:

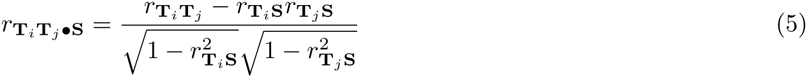

As before, we can rewrite this using *Z*-score formulation. Let us denote the adjusted transcriptional profile of tissue *i* as **Y**_*i*_ = **Z**_*i*_ – *r*_**T*i*S**_**S**. We define the *adjusted transcriptional signature* of tissue *i* as 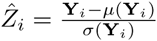. Finally, we can rewrite the adjusted transcriptional similarity as:

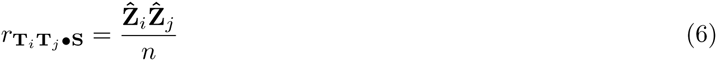

We use this approach, referred to as Peeling the Common Signature (PCS) hereafter, to remove the shared subspace of a given set of expression profiles and construct the corresponding adjusted transcriptional signatures. Significantly positive transcriptional similarities in this framework are indicators of shared tissue-specific pathways, whereas negative scores are indicators of *mutually exclusive* pathways that are used by one tissue but are absent in the other. We use these adjusted transcriptional signatures in our study to identify marker genes. However, when applying methods that rely on the positivity of input expression matrix, such as the entropy-based method by Schug *et al.*[32], one can use the sigmoid transform of these scores as follows:

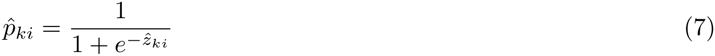

Please note that this transformation, when applied to the raw transcriptional signatures, is equivalent to the previously known softmax normalization:

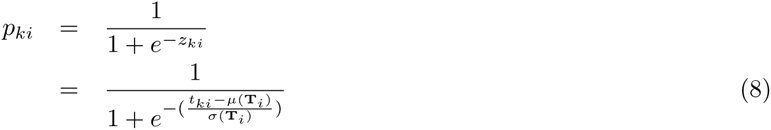

This normalization is known to remove the effect of outliers, while preserving a linear relationship for the mid-range values. We emphasize here that our proposed PCS approach is universally applicable to different marker detection methods, and can be used as an independent pre-processing step.

### Identifying novel marker genes in the reduced subspace of adjusted signatures

Different methods for identifying marker genes not only differ in their underlying formulation, but also make different assumptions regarding the nature of tissue-selective genes:

#### Restricted expression domain

Marker genes are assumed to be expressed in only a limited number of tissues/ cell-types. Genes that are universally expressed in a large number of tissues are less-informative with respect to uncovering tissue-specific functions.

#### Reproducibility

Marker genes should exhibit low variability, both within the cell-type(s) of interest, as well as in the tissues in which they are not expressed. This is a critical assumption, which ensures the reproducibility of the results, since genes that are time or microenvironment dependent are not only poor markers for cell-types, but can change from experiment to experiment.

#### High relative expression

Marker genes are assumed to have much higher expression in the host tissue/ cell-type, compared to the rest of tissues. This is a common assumption, especially in cases where biological replicas are either not available, or there are only a few of them. The significant gap between the average expression of marker genes in the host cell versus unrelated cell-types ensures that even with nominal level of biological variability, these markers are still distinguishable.

#### High absolute expression

Marker genes should have a globally high expression level.

The last assumption is of significance, specially for markers that are to be used for cell-sorting or drug delivery to specific cells, as in the new antibody-conjugated drugs being developed by major vendors. However, this is typically ignored because of experimental limitations, since the absolute value of expression is not comparable across genes because of the experimental biases. We propose a novel method for identifying genes, which focuses on the first three modeling assumptions. Our method is applicable to cases in which no biological replicas are available, or their number is not enough to estimate biological variation of genes. We use cross tissue variations as a proxy for estimating this parameter and ensuring reproducibility. Our method uses the adjusted transcriptional signatures as an input. Then, it constructs groups of similar tissues using these signatures, and reduces the space of tissues and genes by repeated application of the PCS method, and representing each group using its signature vector, while keeping ungrouped tissues as they are. Let us denote the sorted expression value of a given gene *k* by a vector **x** of size *q*. We use a *t*-test type method to assess the separability of the expression domain for gene *k*, encoded by vector **x**, and ensure *reproducibility* of the results. For each cut of size *i* in **x**, we calculate an enrichment score as follows:

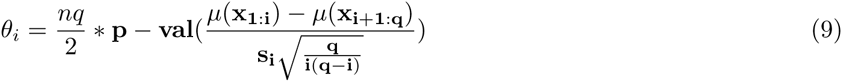

where 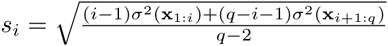 the *p*-value is computed using the Student’s t-distribution with *q* – 2 degrees of freedom, and the 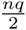 factor corrects for multiple hypothesis testing using the Bonferroni method. To ensure *restriction of the expression domain*, we only look at the first 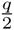 cuts. That is, a gene is declared as preferentially expressed if it is significantly over expressed in a group of at max half of tissues. Using this formulation, we identify the optimal cut as:

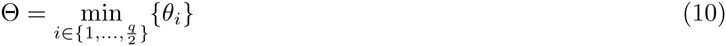

If Θ is greater than a user-provided *p*-value threshold, we will ignore gene *k*. Otherwise, we use the optimal cut to partition tissues into an active set and a null set, denoted by *α* and *η*, respectively. To ensure *high relative expression*, we compute a gap score for gene *k*, represented by *δ*_*k*_, which is defined as the difference between the min of *α* and the max of *η*, normalized by the relative difference of the mean of the two populations, or more formally:

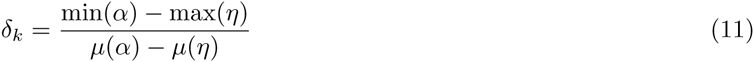

The computed value *δ*_*k*_ is always between 0 and 1, with values closer to 1 being desired. Stated otherwise, *δ*_*k*_ measures the uniformity of gene expression in each group, in a similar fashion to the methods based on entropy. If the mean of one population is significantly different from the marginal values of the cut, then there is a significant difference from the mean estimate of the population, used to assess separability, and the marginal values of the populations. We use a user-provided gap threshold to cut *δ* values and identify genes with high relative expression.

## Author’s contributions

SM proposed the initial idea of research, conceived of the study, implemented the codes, ran the experiments, and prepared the manuscript. AG provided guidance relative to the theoretical and practical aspects of the methods, and design of proper statistical model(s) to validate the results. All authors participated in designing the structure and organization of final manuscript. All authors read and approved the final manuscript.

## Acknowledgements

This work is supported by the Center for Science of Information (CSoI), an NSF Science and Technology Center, under grant agreement CCF-0939370, and by NSF Grant BIO 1124962.

## Additional Files

### Additional file 1 — Raw transcriptional signatures

This is a comma-separated file in *.csv format. First column is the HUGO gene symbols, and first row is the column header. The rest of the table is the computed raw transcriptional signature of the GNF gene atlas dataset.

### Additional file 2 — Adjusted transcriptional signatures

This is a comma-separated file in *.csv format. First column is the HUGO gene symbols, and first row is the column header. The rest of the table is the computed adjusted transcriptional signature of the GNF gene atlas dataset.

### Additional file 3 — Shared markers between monocytes and involuntary muscles

This is a comma-separated file in *.csv format. First column is the HUGO gene symbols for markers, and the second column is the observed difference between involuntary muscles and monocyte (the higher the value, the more significant the expression is in involuntary muscles). There are duplicate genes in column one, which is due to multiple probesets corresponding to the same gene.

### Additional file 4 — Shared functional pathways between monocytes and involuntary muscles

This is an Excel table in *.xls format representing the functional enrichment results using g:Profiler for the monocyte marker genes that are shared with smooth muscle and cardiac myocyte.

### Additional file 5 — Enrichment map of shared markers between monocytes and involuntary muscles

This is a *.pdf file representing the enrichment map of shared markers, constructed using the Gene Enrichment Profiler.

### Additional file 6 — Adjusted transcriptional similarities among all pair of tissues/cell-types

This is a comma-separated file in *.csv format. First and second columns identify the pair of tissues, whereas the third column represents their computed similarity score.

### Additional file 7 — Annotated table of tissue clusters

This is an excel table in *.xls format. Each column represents an identified cluster of similar tissues/ cell-types, and the column headers are descriptive annotations for each cluster.

### Additional file 8 — Adjusted transcriptional signatures reduced over tissue groups

This is a comma-separated file in *.csv format. First column is the HUGO gene symbols, and first row is the column header (name of tissue groups). The rest of the table is the computed adjusted transcriptional signature projected over each group.

### Additional file 9 — Complete list of identified markers

This is an Excel table in *.xls format. Markers corresponding to each of the identified groups are provided in different worksheets.

### Additional file 10 — Complete list of enriched functional terms for the identified brain markers in this study

This is an Excel table in *.xls format representing the enriched functional terms in among brain marker genes, computed using g:Profiler.

### Additional file 11 — Consolidated list of brain-specific markers from previous studies

This is an Excel table in *.xls format. Brain markers from different studies are converted to HUGO gene symbol and are provided in different worksheets.

### Additional file 12 — Enriched disease terms among brain-selective markers

This is an Excel table in *.xls format. First column is the DO ID of the disorder, and the rest of columns indicate the statistical significance of the terms in different brain-selective datasets.

## References

[1] Dezso, Z., Nikolsky, Y., Sviridov, E., Shi, W., Serebriyskaya, T., Dosymbekov, D., Bugrim, A., Rakhmatulin, E., Brennan, R.J., Guryanov, A., Li, K., Blake, J., Samaha, R.R., Nikolskaya, T.: A comprehensive functional analysis of tissue specificity of human gene expression. BMC biology 6, 49 (2008). DOI: 10.1186/1741-7007-6-49

[2] Chang, C.-W., Cheng, W.-C., Chen, C.-R., Shu, W.-Y., Tsai, M.-L., Huang, C.-L., Hsu, I.C.: Identification of human housekeeping genes and tissue-selective genes by microarray meta-analysis. PloS one 6(7), 22859 (2011). DOI: 10.1371/journal.pone.0022859

[3] Souiai, O., Becker, E., Prieto, C., Benkahla, A., De las Rivas J., Brun, C.: Functional integrative levels in the human interactome recapitulate organ organization. PloS one 6(7), 22051 (2011). DOI: 10.1371/journal.pone.0022051

[4] Wang, L., Srivastava, A.K., Schwartz, C.E.: Microarray data integration for genome-wide analysis of human tissue-selective gene expression. BMC genomics 11 Suppl 2, 15 (2010). DOI: 10.1186/1471-2164-11-S2-S15

[5] Cavalli, F.M.G., Bourgon, R., Huber, W., Vaquerizas, J.M., Luscombe, N.M.: SpeCond: a method to detect condition-specific gene expression. Genome biology 12(10), 101 (2011). DOI: 10.1186/gb-2011-12-10-r101

[6] Teng, S., Yang, J.Y., Wang, L.: Genome-wide prediction and analysis of human tissue-selective genes using microarray expression data. BMC medical genomics 6 Suppl 1, 10 (2013). DOI: 10.1186/1755-8794-6-S1-S10

[7] Goh, K.-I., Cusick, M.E., Valle, D., Childs, B., Vidal, M., Barabási, A.-L.: The human disease network. Proceedings of the National Academy of Sciences of the United States of America 104(21), 8685–90 (2007). DOI: 10.1073/pnas.0701361104

[8] Lage, K., Hansen, N.T., Karlberg, E.O., Eklund, A.C., Roque, F.S., Donahoe, P.K., Szallasi, Z., Jensen, T.S.t., Brunak, S.r.: A large-scale analysis of tissue-specific pathology and gene expression of human disease genes and complexes. Proceedings of the National Academy of Sciences of the United States of America 105(52), 20870–5 (2008). DOI: 10.1073/pnas.0810772105

[9] Kadota, K., Ye, J., Nakai, Y., Terada, T., Shimizu, K.: ROKU: a novel method for identification of tissue-specific genes. BMC bioinformatics 7, 294 (2006). DOI: 10.1186/1471-2105-7-294

[10] Cheng, W.-C., Tsai, M.-L., Chang, C.-W., Huang, C.-L., Chen, C.-R., Shu, W.-Y., Lee, Y.-S., Wang, T.-H., Hong, J.-H., Li, C.-Y., Hsu, I.C.: Microarray meta-analysis database (M(2)DB): a uniformly pre-processed, quality controlled, and manually curated human clinical microarray database. BMC bioinformatics 11, 421 (2010). DOI: 10.1186/1471-2105-11-421

[11] Uhlén, M., Fagerberg, L., Hallström, B.M., Lindskog, C., Oksvold, P., Mardinoglu, A., Sivertsson, A.s., Kampf, C., Sjöstedt, E., Asplund, A., Olsson, I., Edlund, K., Lundberg, E., Navani, S., Szigyarto, C.A.-k., Odeberg, J., Djureinovic, D., Takanen, J.O., Hober, S., Alm, T., Edqvist, P.-h., Berling, H., Tegel, H., Mulder, J., Rockberg, J., Nilsson, P., Schwenk, J.M., Hamsten, M., Feilitzen, K.V., Forsberg, M., Persson, L., Johansson, F., Zwahlen, M., Heijne, G.V., Nielsen, J., Pontén, F.: Tissue-based map of the human proteome (2015). DOI: 10.1126/science.1260419

[12] Eisenberg, E., Levanon, E.Y.: Human housekeeping genes, revisited (2013). DOI: 10.1016/j.tig.2013.05.010

[13] Yanai, I., Benjamin, H., Shmoish, M., Chalifa-Caspi, V., Shklar, M., Ophir, R., Bar-Even, A., Horn-Saban, S., Safran, M., Domany, E., Lancet, D., Shmueli, O.: Genome-wide midrange transcription profiles reveal expression level relationships in human tissue specification. Bioinformatics (Oxford, England) 21(5), 650–9 (2005). DOI: 10.1093/bioinformatics/bti042

[14] Barber, R.D., Harmer, D.W., Coleman, R.A., Clark, B.J.: GAPDH as a housekeeping gene: analysis of GAPDH mRNA expression in a panel of 72 human tissues. Physiological genomics 21(3), 389–95 (2005). DOI: 10.1152/physiolgenomics.00025.2005

[15] Abbas, A.R., Baldwin, D., Ma, Y., Ouyang, W., Gurney, A., Martin, F., Fong, S., van Lookeren Campagne M., Godowski, P., Williams, P.M., Chan, A.C., Clark, H.F.: Immune response in silico (IRIS): immune-specific genes identified from a compendium of microarray expression data. Genes and immunity 6(4), 319–31 (2005). DOI: 10.1038/sj.gene.6364173

[16] McKenna, K., Beignon, A.-S., Bhardwaj, N.: Plasmacytoid dendritic cells: linking innate and adaptive immunity. Journal of virology 79(1), 17–27 (2005). DOI: 10.1128/JVI.79.1.17-27.2005

[17] Zuccolo, J., Unruh, T.L., Deans, J.P.: Efficient isolation of highly purified tonsil B lymphocytes using Rosette-Sep with allogeneic human red blood cells. BMC immunology 10, 30 (2009). DOI: 10.1186/1471-2172-10-30

[18] Reimand, J., Arak, T., Vilo, J.: g: Profiler–a web server for functional interpretation of gene lists (2011 update). Nucleic acids research 39(Web Server issue), 307–15 (2011). DOI: 10.1093/nar/gkr378

[19] Benita, Y., Cao, Z., Giallourakis, C., Li, C., Gardet, A., Xavier, R.J.: Gene enrichment profiles reveal T-cell development, differentiation, and lineage-specific transcription factors including ZBTB25 as a novel NF-AT repressor. Blood 115(26), 5376–84 (2010). DOI: 10.1182/blood-2010-01-263855

[20] Shea-Donohue, T., Notari, L., Sun, R., Zhao, A.: Mechanisms of smooth muscle responses to inflammation (2012). DOI: 10.1111/j.1365-2982.2012.01986.x

[21] Gerthoffer, W.T.: Mechanisms of vascular smooth muscle cell migration (2007). DOI: 10.1161/01.RES.0000258492.96097.47

[22] Nelson, P.R., Yamamura, S., Mureebe, L., Itoh, H., Kent, K.C.: Smooth muscle cell migration and proliferation are mediated by distinct phases of activation of the intracellular messenger mitogen-activated protein kinase. Journal of Vascular Surgery 27(1), 117–125 (1998). DOI: 10.1016/S0741-5214(98)70298-8

[23] Hayes, I.M., Jordan, N.J., Towers, S., Smith, G., Paterson, J.R., Earnshaw, J.J., Roach, A.G., Westwick, J., Williams, R.J.: Human vascular smooth muscle cells express receptors for CC chemokines. Arteriosclerosis, thrombosis, and vascular biology 18(3), 397–403 (1998). DOI: 10.1161/01.ATV.18.3.397

[24] Halwani, R., Al-Abri, J., Beland, M., Al-Jahdali, H., Halayko, A.J., Lee, T.H., Al-Muhsen, S., Hamid, Q.: CC and CXC chemokines induce airway smooth muscle proliferation and survival. Journal of immunology (Baltimore, Md.: 1950) 186(7), 4156–4163 (2011). DOI: 10.4049/jimmunol.1001210

[25] Yun Kui Zhu, Liu, X., Wang, H., Kohyama, T., Wen, F.Q., Sköld, C.M., Rennard, S.I.: Interactions between monocytes and smooth-muscle cells can lead to extracellular matrix degradation. Journal of Allergy and Clinical Immunology 108(6), 989–996 (2001). DOI: 10.1067/mai.2001.120193

[26] Karsdal, M.a., Nielsen, M.J., Sand, J.M., Henriksen, K., Genovese, F., Bay-Jensen, A.-C., Smith, V., Adamkewicz, J.I., Christiansen, C., Leeming, D.J.: Extracellular matrix remodeling: the common denominator in connective tissue diseases. Possibilities for evaluation and current understanding of the matrix as more than a passive architecture, but a key player in tissue failure. Assay and drug development technologies 11(2), 70–92 (2013). DOI: 10.1089/adt.2012.474

[27] Verma, R.P., Hansch, C.: Matrix metalloproteinases (MMPs): Chemical-biological functions and (Q)SARs (2007). DOI: 10.1016/j.bmc.2007.01.011

[28] Reverter, A., Chan, E.K.F.: Combining partial correlation and an information theory approach to the reversed engineering of gene co-expression networks. Bioinformatics (Oxford, England) 24(21), 2491–7 (2008). DOI: 10.1093/bioinformatics/btn482

[29] Su, G., Morris, J.H., Demchak, B., Bader, G.D.: Biological Network Exploration with Cytoscape 3. John Wiley & Sons, Inc., ??? (2002). DOI: 10.1002/0471250953.bi0813s47. http://dx.doi.org/10.1002/0471250953.bi0813s47

[30] Uhlen, M., Oksvold, P., Fagerberg, L., Lundberg, E., Jonasson, K., Forsberg, M., Zwahlen, M., Kampf, C., Wester, K., Hober, S., Wernerus, H., Björling, L., Ponten, F.: Towards a knowledge-based Human Protein Atlas. (2010). DOI: 10.1038/nbt1210-1248

[31] Kasprzyk, A.: BioMart: driving a paradigm change in biological data management. Database: the journal of biological databases and curation 2011, 049 (2011). DOI: 10.1093/database/bar049

[32] Schug, J., Schuller, W.-P., Kappen, C., Salbaum, J.M., Bucan, M., Stoeckert, C.J.: Promoter features related to tissue specificity as measured by Shannon entropy. Genome biology 6(4), 33 (2005). DOI: 10.1186/gb-2005-6- 4-r33

[33] Yu, G., Wang, L.-G., Yan, G.-R., He, Q.-Y.: DOSE: an R/Bioconductor package for disease ontology semantic and enrichment analysis. Bioinformatics 31(4), 608–609 (2014). DOI: 10.1093/bioinformatics/btu684

[34] Kibbe, W.A., Arze, C., Felix, V., Mitraka, E., Bolton, E., Fu, G., Mungall, C.J., Binder, J.X., Malone, J., Vasant, D., Parkinson, H., Schriml, L.M.: Disease Ontology 2015 update: an expanded and updated database of human diseases for linking biomedical knowledge through disease data. Nucleic Acids Research 43(D1), 1071–1078 (2014). DOI: 10.1093/nar/gku1011

[35] Su, A.I., Wiltshire, T., Batalov, S., Lapp, H., Ching, K.A., Block, D., Zhang, J., Soden, R., Hayakawa, M., Kreiman, G., Cooke, M.P., Walker, J.R., Hogenesch, J.B.: A gene atlas of the mouse and human protein-encoding transcriptomes. Proceedings of the National Academy of Sciences of the United States of America 101(16), 6062–7 (2004). DOI: 10.1073/pnas.0400782101

[36] Wu, C., Macleod, I., Su, A.I.: BioGPS and MyGene.info: organizing online, gene-centric information. Nucleic Acids research 41(Database issue), 561–5 (2013). DOI: 10.1093/nar/gks1114

[37] Wu, C., Orozco, C., Boyer, J., Leglise, M., Goodale, J., Batalov, S., Hodge, C.L., Haase, J., Janes, J., Huss, J.W., Su, A.I.: BioGPS: an extensible and customizable portal for querying and organizing gene annotation resources. Genome biology 10(11), 130 (2009). DOI: 10.1186/gb-2009-10-11-r130

